# Phospholipid exchange shows insulin receptor activity is supported by both the propensity to form wide bilayers and ordered domains (rafts)

**DOI:** 10.1101/2021.04.27.440802

**Authors:** Pavana Suresh, W. Todd Miller, Erwin London

## Abstract

Using efficient methyl-alpha-cyclodextrin mediated lipid exchange, we studied the effect of altering plasma membrane outer leaflet phospholipid composition upon the activity of insulin receptor (IR) in mammalian cells. After substitution of endogenous lipids with lipids having an ability to form liquid ordered (Lo) domains (sphingomyelins) or liquid disordered (Ld) domains (unsaturated phosphatidylcholines (PCs)), we found that the propensity of lipids to form ordered domains is required for high IR activity. Additional substitution experiments using a series of saturated PCs showed that IR activity increased substantially with increasing acyl chain length. Increasing acyl chain length increases both bilayer width and the propensity to form ordered domains. To distinguish the effects of membrane width and domain formation, we incorporated purified IR into alkyl maltoside micelles with increasing hydrocarbon lengths. IR activity increases with increased chain length, but more modestly than by increasing lipid acyl chain length in cells. This suggests that the ability to form Lo domains as well as wide bilayer width contributes to increased IR activity. Inhibition of phosphatases with sodium orthovanadate showed that some of the lipid dependence of IR activity upon lipid structure reflected protection from phosphatases by lipids that support Lo domain formation. The results are consistent with a model in which a combination of bilayer width and ordered domain formation modulate IR activity via effects upon IR conformation and accessibility to phosphatases.

**Significance:** This study shows how methyl-α-cyclodextrin mediated lipid exchange can be used to probe the influence of lipid structure upon the functioning of a transmembrane receptor. Plasma membranes having a propensity to form Lo domains are required to support a high level of IR activity. The studies indicate this may reflect an effect of lipid environment upon IR domain localization, which in turn alters its conformation and vulnerability to phosphatases. Alterations in lipid composition could conceivably regulate IR activity *in vivo*.

## INTRODUCTION

Signal transduction is thought to be influenced by lipid organization into domains that can facilitate receptor clustering [1]. The two types of domains thought to exist in cells are liquid ordered domains (Lo) also known as lipid rafts, and liquid disordered domains (Ld) [2]. Lipid rafts are predominantly composed of saturated lipids such as sphingomyelin (SM), and cholesterol, with properties such as tight lipid packing, increased membrane thickness, and slower lateral diffusion rates relative to Ld domains, which are enriched in unsaturated phospholipids [3]. Rafts have been associated in signaling by membrane growth factor receptors [4], immunoglobulin E receptors, and antigen receptors in T and B cells [5]–[7]. Previous studies have reported that some receptors have preferential distribution in lipid rafts and/or that their activity responds to cellular cholesterol levels, which can alter domain formation and properties [4], [8].

Insulin receptor (IR) is a receptor tyrosine kinase involved in cellular glucose intake and lipid metabolism [9], [10].There is evidence associating IR activity with lipid rafts [11]. Studies have reported a decrease in IR autophosphorylation activity, reduction in downstream signaling protein IRS-1, and a reduction in glucose uptake by cells after removal of raft-promoting cholesterol from cells [12]–[14]. However, cholesterol depletion has many effects upon membrane structure, and cannot by itself demonstrate a functional role for lipid rafts [15]–[17]. IR has also been shown to be sequestered in detergent resistant membrane fractions after insulin stimulation [18], and in the related caveolae fractions after insulin stimulation [12], [13], [19]. However, the relationship of detergent resistant membranes and caveolae to Lo domains in cells is complex [6], [20], [21]. Thus, these observations are insufficient to definitively demonstrate an interaction of IR with lipid rafts in vivo, or that lipid rafts have a role in IR function [2], [22].

By using methyl-β-cyclodextrin (MβCD) to substitute cellular cholesterol with sterols having various propensities to aid Lo domain formation, we previously reported that IR autophosphorylation activity was only high in cells containing sterols having a propensity to form ordered domains [23]. This modulation of IR activity could reflect one of several effects of sterol substitutions: altered ordered domain formation, altered membrane width, or altered lipid packing in a homogenous membrane lacking domains. To investigate further, we performed outer leaflet lipid exchange on CHO cells stably expressing IR (CHO IR), replacing endogenous plasma membrane outer leaflet lipids with a series of phospholipids (including sphingomyelin) that support or disrupt rafts. We found that phospholipids with a propensity to form Lo domains resulted in activation of IR. This effect was partly due to the decreased accessibility of IR to phosphatases after substitution using phospholipids that promote Lo domain formation. In addition, we found that increased hydrocarbon width promoted IR activity. A model is proposed for how association of IR with Lo domains could control its activity.

## METHODS

### Materials

Studies used previously generated Chinese hamster ovary cells stably expressing human IR (CHO IR cells) [24]. 293T cells stably expressing human IR (293T IR cells) with a C terminal streptavidin-binding peptide (SBP) tag were generated previously [23], [25]. Brain sphingomyelin (bSM), egg sphingomyelin (eSM), 1-palmitoyl-2-oleoyl-sn-glycerol-3-phosphocholine (POPC), 1,2-dioleoyl-sn-glycero-3-phosphocholine (DOPC), 1,2-dilauroyl-sn-glycero-3-phosphocholine (DLPC), 1,2-dimyristoyl-sn-glycero-3-phosphocholine (DMPC), 1,2-dipalmitoyl-sn-glycero-3-phosphocholine (DPPC), 1,2-distearoyl-sn-glycero-3-phosphocholine (DSPC) and cholesterol were from Avanti Polar Lipids (Alabaster, AL). Methyl-α-cyclodextrin (MαCD) was purchased from AraChem (Budel, the Netherlands). Methotrexate and sodium orthovanadate (SOV) were purchased from Sigma Aldrich (St. Louis, MO). Dulbecco’s modified eagle medium (DMEM, 4.5g/L glucose, L-glutamine, sodium pyruvate), phosphate buffered saline (PBS) without calcium and magnesium (0.144 g/L KH_2_PO_4_, 9 g/L NaCl, 0.795 g/L Na_2_HPO_4_ (anhydrous)), trypsin-EDTA, antibiotic-antimycotic solution, L-glutamine were purchased from Corning (Corning, NY). Non-essential amino acids and ham’s F12 media were purchased from Gibco. G418 (geneticin) was from Goldbio (St Louis, MO), fetal bovine serum (FBS) was from VWR international (Radnor, PA), n-tetradecyl-β-D-maltopyranoside (TDM) and n-hexadecyl-β-D-maltopyranoside (HDM) were from Anatrace (Maumee, OH). N-decyl-β-D-maltopyranoside (DM) from Dojindo Molecular Technologies (Rockville, MD), n-dodecyl-β-D-maltopyranoside (DDM) from Thermo Fisher (Waltham, MA).

### Lipid and cholesterol purity

Lipid stock solutions in chloroform were checked for purity by chromatographing them on a high performance thin layer chromatography (HP-TLC) in 65:35:5 (v:v:v) chloroform:methanol:28.0-30.0 (v/v) % ammonium hydroxide solvent. All lipids were present as single bands except eSM and bSM which have a mixture of acyl chain lengths and thus show multiple bands. Cholesterol stock solution was similarly checked for purity in 3:2 (v:v) hexanes:ethyl acetate solvent and a single band was observed.

### Cell culture

CHO IR cells were grown in DMEM supplemented with 10% FBS, 300 µg/ml glutamine, 100 µg/ml non-essential amino acids, 50 µg/ml G418, 2 µM methotrexate, and antibiotic-antimycotic solution (diluting 100X stock into medium). 293T IR cells were cultured in DMEM 4.5 g/L glucose media supplemented with 10% FBS and antibiotic-antimycotic solution. All cells were grown in a 37 °C incubator with 5% CO_2_. CHO IR cells were starved in serum free ham’s F12 media.

### Preparation of lipid exchange media

Lipid exchange media with lipid-loaded MαCD was prepared as described previously [26], [27]. Briefly, the desired amounts of lipids in chloroform were dried under N_2_ gas stream and further dried under high vacuum for 1 h. Multilamellar vesicles (MLVs) were formed by adding serum free ham’s F12 media to dried lipids and incubating in a 70 °C water bath for 5 min followed by vortexing to disperse the lipid. MLVs and MαCD (from a stock of 300-400 mM in PBS) were added to serum free ham’s F12 media to prepare lipid exchange mixtures with the desired final concentrations. These were solutions with 40 mM MαCD and 0.5 mM lipid for DSPC, 1 mM lipid for bSM, eSM, and DPPC, 2 mM lipid for DMPC, and 4 mM lipid for POPC, DOPC, and DLPC. All lipid exchange mixtures except for DPPC and DSPC were incubated in 37 °C water bath for 30 min to allow lipids to be loaded onto MαCD and incubated in room temperature for 20 min before adding to cells. DPPC and DSPC exchange mixtures were incubated in 70 °C water bath for 30 min and then placed in room temperature for 30 min before adding to cells.

### Lipid exchange in CHO IR cells

Cells were grown to 80-90% confluency in 60 mm plates and lipid exchange was performed as described [27]. Briefly, cells were washed in PBS twice at room temperature and starved in serum free media for 22 h at 37 °C in a 5% CO_2_ incubator. The cells were then washed in PBS once and 1 ml lipid exchange mixture was added. Exchange was carried out at 26-27 °C for 1 h in an incubator without CO_2_. After exchange, cells were washed in PBS thrice.

### IR autophosphorylation

After lipid exchange, CHO IR cells in 60 mm plates were incubated with or without 100 nM insulin (in serum free ham’s F-12 media) for 5 min at room temperature. Stimulation was terminated by removing insulin-containing media and adding 1 ml cold PBS after a 1 ml cold PBS wash. Cells were scraped from the plate and pelleted by centrifugation for 5 min at 1000 x g (at 4 °C) and lysed with 150-200 µl lysis buffer (50 mM Tris pH 8.0, 200 mM NaCl, 1% (v/v) Triton X-100, 1 mM EDTA, 1% (w/v) sodium deoxycholate, 10 µg/ml leupeptin, 10 µg/ml aprotinin and 1 mM activated sodium orthovanadate). Lysates were centrifuged at 16,837 x g for 10 min (at 4 °C), aliquots were removed and then mixed with 5x Laemmli buffer [28] for western blotting analysis of autophosphorylation and protein levels. The remaining lysates were used to determine protein concentration using the Bradford assay [29].

For measurement of IR autophosphorylation at 37°C after 1 h of lipid exchange in 60 mm plates, cells were washed in warm PBS (pre-warmed at 37°C) thrice. Then 100 nM insulin in serum free ham’s F-12 media (pre-warmed at 37°C) was added, and cells were incubated in 37°C, 5% CO_2_ incubator for 5 min. After insulin stimulation, insulin was removed, and cells were washed once in warm PBS (pre-warmed at 37°C). Lysis buffer was added directly on plate and cell lysates were collected by scraping. Lysates were centrifuged at 16,837 x g for 10 min (at RT), aliquots were removed and then mixed with 5x Laemmli buffer for western blotting analysis of autophosphorylation and protein levels. The remaining lysates were used to determine protein concentration using the Bradford assay.

### Western blotting

Whole cell lysate samples (typically 5-10 µg total protein) were loaded in equal amounts across all lanes (based on protein concentration using Bradford assay) and run on 7.5 % acrylamide SDS-PAGE and then transferred to polyvinylidene fluoride (PVDF) membranes (Millipore, Burlington, MA) for 1 h at 100 V at 4°C. Membranes were blocked with 5% (w/v) BSA in Tris-Buffered Saline and Tween 20 (TBST, 20 mM Tris, 137 mM NaCl, and 0.1 % v/v Tween 20, pH 7.6) for 1 h and then incubated with primary antibodies overnight at 4°C, followed by secondary antibodies for 30 min at room temperature. Membranes were imaged using western blotting substrate (Thermo scientific pierce ECL, Waltham, MA) and exposed to autoradiographic film (Bioexcell, Lebanon, NH). Primary antibodies used were Anti-pYpY1162/1163 IR (Catalog number AF2507, R&D systems Inc., Minneapolis, MN), anti-insulin receptor β (Catalogue number 3025, Cell signaling technology, Danvers, MA), and anti-phosphotyrosine (Catalogue number 05-321, Millipore, Burlington, MA). Secondary antibodies used were rabbit IgG HRP conjugated (GE Healthcare life sciences, Malborough, MA) or mouse IgG HRP conjugated (GE Healthcare life sciences, Malborough, MA). All primary antibodies were diluted 1:1000 in TBST with 5% BSA from the commercially provided solution, except pYpY1162/1163 IR which was used at 1:400 dilution in TBST with 5% BSA. Secondary antibodies were diluted to 1:5000 in TBST. To avoid having to strip membranes duplicate gels were run, one for probing with pYpY1162/1163 IR and one for IR-β.

### Sodium orthovanadate (SOV) treatment with lipid exchange

Lipid exchange was carried out as described (see above) with activated SOV [30] added prior to exchange to a concentration of 1 mM in 1 ml of exchange media. For untreated samples SOV was added at a concentration of 1 mM in 1 ml serum free ham’s F12.

### Insulin receptor purification from 293T IR cells for maltoside detergent assay

293T IR cells (at 95% confluency) were harvested from eight 15 cm plates in PBS by pipetting. The cells from the plates were combined and centrifuged at 1000 x g for 5 min at 4 °C. Supernatant was removed and cell pellet was stored at -80 °C. For IR purification the pellet was lysed at 4 °C with 40 ml purification lysis buffer (20 mM Tris pH 8.0, 400 mM NaCl, 10% (v/v) glycerol, 1% (v/v) Triton X-100, 1 mM EDTA, 5 µg/ml leupeptin, 5 µg/ml aprotinin) while mixing on an end-over-end rotor for 1 h. All the following steps were also at 4°C unless otherwise noted. Lysate was then centrifuged at 12,520 x g for 30 min and the supernatant solution was filtered through a 0.8 µm syringe filter (Millex-AA, Millipore, Burlington, MA) before adding it to 2 ml of Strep-Tactin superflow resin (Qiagen, Hilden, Germany) which was pre-equilibrated with 10 ml lysis buffer. Cleared lysate was incubated with resin for 30 min before pouring it into a 1.5 cm diameter glass chromatography column. The column was then rinsed with 30 column volumes (CV) of buffer A (20 mM Tris pH 8.0, 200 mM NaCl, 10% glycerol, 0.1 % (v/v) Triton X-100). Beads in 50% slurry were transferred to a 50 ml polypropylene conical tube (Corning, Corning, NY) and basal phosphorylation was removed by treatment with soluble glutathione s-transferase tagged *Yersinia* tyrosine phosphatase (YOP) for 30 min at room temperature on an end-to-end rotor. YOP phosphatase was inhibited with 20 mM activated SOV [30] and incubated for an additional 10 min at room temperature. Beads were transferred back into the column and soluble YOP was washed away with 30 CV of 0.1% Triton X-100 wash buffer. The detergent was then changed to DDM by washing the beads with 10 CV of Buffer B (20 mM Tris pH 8.0, 200 mM NaCl, 0.29 mM DDM). IR was eluted in buffer B containing 2.5 mM D-desthiobiotin. Purified IR was used without an additional concentration step.

### In-vitro autophosphorylation in alkyl maltoside micelles

IR eluted in DDM was diluted 1:10 into solutions containing DM, DDM, TDM or HDM at a concentration 1.25 times (0.5 + critical micelle concentration (CMC)) mM detergent in buffer C (20 mM Tris pH 8.0, 200 mM NaCl). For each detergent condition, 40 µl of IR in each alkyl maltoside detergent was added to 10 µl kinase reaction buffer (1 mM ATP, 1 mM MgCl_2_, 1 mM activated SOV in Tris pH 8) with or without 100 nM insulin to start the reaction and incubated for 5 min in a 23 °C water bath. The final detergent concentration was (0.5 + CMC) mM (i.e. for DDM with CMC 0.17 mM, the final concentration was 0.5 mM + 0.17 mM = 0.67 mM). The reactions were stopped by adding 15 µl of 5x Laemmli buffer and analyzed by western blotting.

### In-vitro radiometric kinase assay

IR in alkyl maltoside detergents were prepared as for autophosphorylation reactions to give (0.5 +CMC) mM final detergent concentration in kinase reactions. Reactions were started by adding 40 µl IR in alkyl maltoside micelles to 10 µl radioactive kinase reaction buffer (0.4 mM ATP, 1 µCi [γ-^32^P]-ATP (PerkinElmer, 10 mCi/mL, 25-50 cpm/mol), 1 mM MgCl_2_, 0.7 mM synthetic peptide KKEEEEYMMMMG (E_4_YM_4_), 1 mM SOV, and 1 mM BSA) with or without 100 nM insulin. The reactions were incubated in 30 °C water bath for 15 min and quenched by adding 18 µl cold 50% (v/v) trichloroacetic acid. Samples were centrifuged at 9,296 x g for 2 min to pellet IR and 35 µl of supernatant was spotted on P81 phospho-cellulose paper (Whatman). Remaining [γ-^32^P]-ATP was removed by washing P81 papers in 0.5% (v/v) cold phosphoric acid thrice for 10 min each. Finally, P81 papers were dried, and radioactivity was measured in Hidex 300 SL scintillation counter.

### Lipid extraction from cells

For lipid exchange samples, cells were air dried on plate after PBS washes for 10 min in room temperature and then 1 ml of 3:2 (v:v) hexanes:isopropanol was added to each plate. The plates were kept on a rocker for 30 min and the organic solvent was transferred to a new borosilicate glass test tube and stored in -20 °C. The remaining cell debris on plate was dissolved in 1 N NaOH and used for protein quantification by Bradford assay.

### HP-TLC of lipids

Extracted lipids were dried under N_2_ gas and then re-dissolved in 1:1 (v:v) chloroform:methanol. Aliquots were loaded on HP-TLC plates (HP-TLC Silica Gel 60 plates (Merck, Kenilworth, NJ)). Protein quantification showed samples should have similar amounts of lipid loaded per lane. Lipid samples were chromatographed in 65:35:5 (v:v:v) chloroform:methanol:28.0-30.0 (v/v) % ammonium hydroxide to separate phospholipids. For measuring lipid content, plates were sprayed with a solution of 3% (w/v) cupric acetate and 8% (v/v) phosphoric acid dissolved in water. The plates were air dried and charred at 180-200 °C to detect lipid bands. Band intensity was then measured using ImageJ program. For lipid exchange samples, exogenous lipid introduced was estimated from the decrease in cellular SM bands, as staining is near linear in SM concentration [31].

### Flow cytometry for detecting IR on plasma membrane

After lipid exchange as described above, cells were dislodged from plates using enzyme free cell dissociation solution (Millipore, Burlington, MA), spun down at 300 x g, 4 °C for 5 min and washed with blocking buffer (0.5% BSA, 2 mM EDTA in PBS) at room temperature. Manufacturer’s protocol was followed for fluorescent antibody staining. Cells were resuspended in 100 µl blocking buffer, CD220-PE antibodies against IR alpha chain or recombinant antibody (REA) control (S)-PE (Miltenyi Biotech, Bergisch Gladbach, Germany) were added, and the sample incubated at 4 °C in the dark for 10 min. Samples were washed at room temperature in PBS blocking buffer again. Flow cytometry analysis was performed using FACSCalibur (BD Bioscience, San Jose, CA) collecting 10,000 events per sample. Data was analyzed using FlowJo version 10 software.

### Cell viability assessment using flow cytometry

After lipid exchange and an additional 1 h recovery step in complete media in a 37°C, 5% CO_2_ incubator, cell viability was assessed using propidium iodide (Thermo Fisher, Waltham, MA) as described previously [27]. Briefly, after lipid exchange, cells were washed in PBS thrice and 1 ml complete media was added. The cells were then incubated in a 37°C, 5% CO_2_ incubator for 1 h which was followed by a PBS wash. The cells were then detached using trypsin, washed in PBS, and resuspended in 100 µl of binding buffer (50 mM HEPES pH 7.4, 700 mM NaCl, 12.6 mM CaCl_2_) per million cells. Propidium iodide was added (1 μg/ml) and samples were incubated at room temperature while protected from light for 15 min on a shaker. Samples were diluted with 400 µl binding buffer and analyzed using FACSCalibur (BD Biosciences, San Hose, CA) flow cytometer collecting 50,000 events per sample.

### Quantification of western blot images and ensuring linearity of detection

Western blot bands were quantified using ImageJ software analysis. Individual pYpY IR autophosphorylation bands were normalized to total IR-β bands for corresponding samples. To confirm the signal from antibodies is linear in concentration at the exposure used, a standard curve of one sample (usually untreated sample or sample with the highest intensity) was run on the same blot as the experiments. Averages and standard deviations from independent experiments were then calculated.

## RESULTS

### Lipid exchange is highly efficient and shows unaltered IR surface expression and minimal cell damage

We first carried out phospholipid exchange in CHO IR cells and analyzed the resulting change in SM content. **Figure 1A** shows exchange with SM increased the total SM content and exchange with PC decreased SM content. The efficiency of exchange by PCs was evaluated by comparing the loss of outer leaflet SM band in untreated cells. This can be used because replacement of endogenous lipid by exogenous lipid upon exchange is roughly 1:1 [26], [32]. Exchange with all PCs (except DPPC) showed ∼ 70% decrease of total SM in cells. This is close to the maximal level (∼ 70-80%) of endogenous cellular SM that can be removed by exchange, with the remaining SM likely being in a pool that is inaccessible to exchange (either being present in the inner leaflet of the plasma membrane or internal organelles)[26], [33]. DPPC gave only 50% replacement of cellular SM, indicating somewhat lower efficiency of exchange. Based on this exchange data we can infer that about 70% of total SM in the whole cell resides in the PM outer leaflet. Assuming exchange with exogenous SM is also highly efficient, (80-100%, as estimated in previous studies [26], [27]), the observed 5-fold increase in SM would correspond to the PM lipid being 11-14% SM. This is in good agreement with the composition of SM content previously reported for CHO cell PM [33]. Overall, exchange appears to be highly efficient, with the exogenous lipid forming the preponderance of phospholipid after exchange.

**Figure 1:**
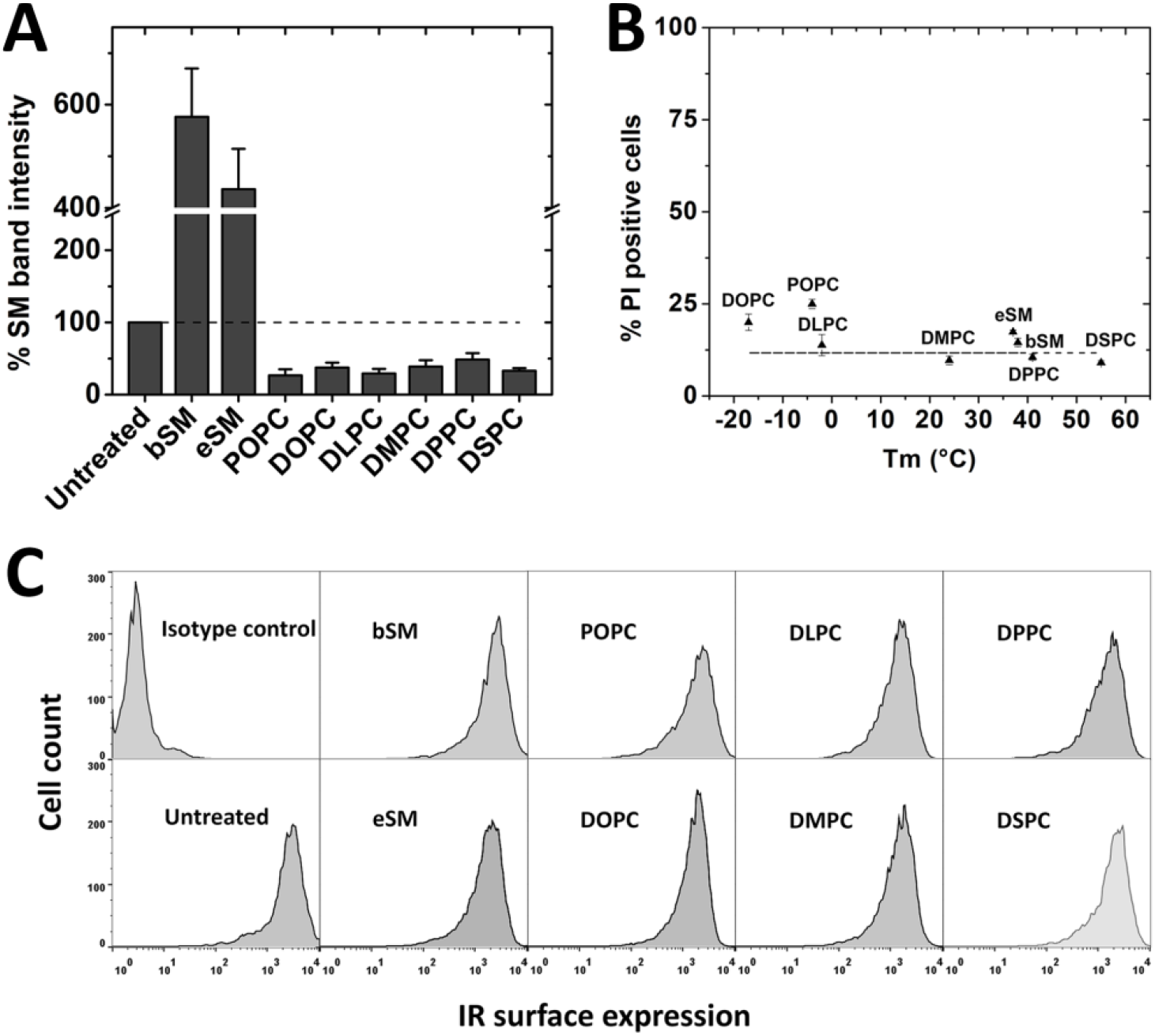
CHO IR exchange efficiency, cell damage and IR surface expression after lipid exchange. (A) Lipid exchange efficiency indicated by change in % SM band intensity compared to untreated cells. The average (mean) SM content ± S.D. from HP-TLC analysis of three independent experiments is shown. A representative HP-TLC image is shown in **supplementary figure 1**. All band intensities are shown normalized to untreated control SM levels set at 100%. A dashed line extends the 100% value for comparison to other lipid exchange samples. (B) Cell damage monitored by propidium iodide (PI) staining using flow cytometry in lipid exchanged samples plotted according to the lipid gel-to-Ld phase transition temperatures (Tm). The average and S.D. from three experiments is shown. (C) Surface expression of IR using phycoerythrin conjugated IR antibody along with isotype control (non-binding antibody).

To ensure the cells were not damaged after lipid exchange, membrane permeability to propidium iodide was monitored using flow cytometry. The measurement was made after a 1 h incubation of lipid-exchanged cells in serum-containing media, as cells can be damaged by immediate processing after lipid exchange [27]. **Figure 1B** shows that cell viability after exchange was similar to that in untreated cells. Similarly, IR surface expression was found to be largely unaltered immediately after lipid exchange as shown in **Figure 1C** by flow cytometry data using anti-IR phycoerythrin conjugated antibody raised against the IR ectodomain.

### Lipid exchange on CHO IR cells show IR activity is supported in lipids that support ordered domains

IR autophosphorylation activity was previously shown to be supported only by sterols that have a high propensity to form ordered domains in 293T IR cells [23]. Similarly, in CHO IR cells we found that cholesterol depletion using MβCD decreased IR autophosphorylation activity and adding cholesterol back restored activity (data not shown). To further explore the effect of lipid structure on IR activity and its possible connection to Lo domain formation, we measured activity after outer leaflet plasma membrane lipid exchange in CHO IR cells with two different sphingomyelins (brain SM (bSM) and egg SM (eSM)), which have a propensity to form Lo domains, and with two unsaturated PCs (1-palmitoyl-2-oleoyl-glycero-3-phosphocholine (POPC) and 1,2-dioleoyl-sn-glycero-3-phosphocholine (DOPC)) which promote formation of the Ld state [34], [35] **[Figure 2]. Figure 2A** and **2B** illustrate representative blots showing the effect of lipid exchange upon IR activity. To confirm western blot bands were within the linear range of detection, a standard curve of signal intensity vs. protein was loaded on each blot **[Figure 2A, 2B and Supplementary Figure 2]**. Quantification of western blot data is shown in **Figure 2C and D**. In subsequent figures, only the analyzed data is presented, but representative western blot images are shown in **Supplementary Figure 3**.

**Figure 2:**
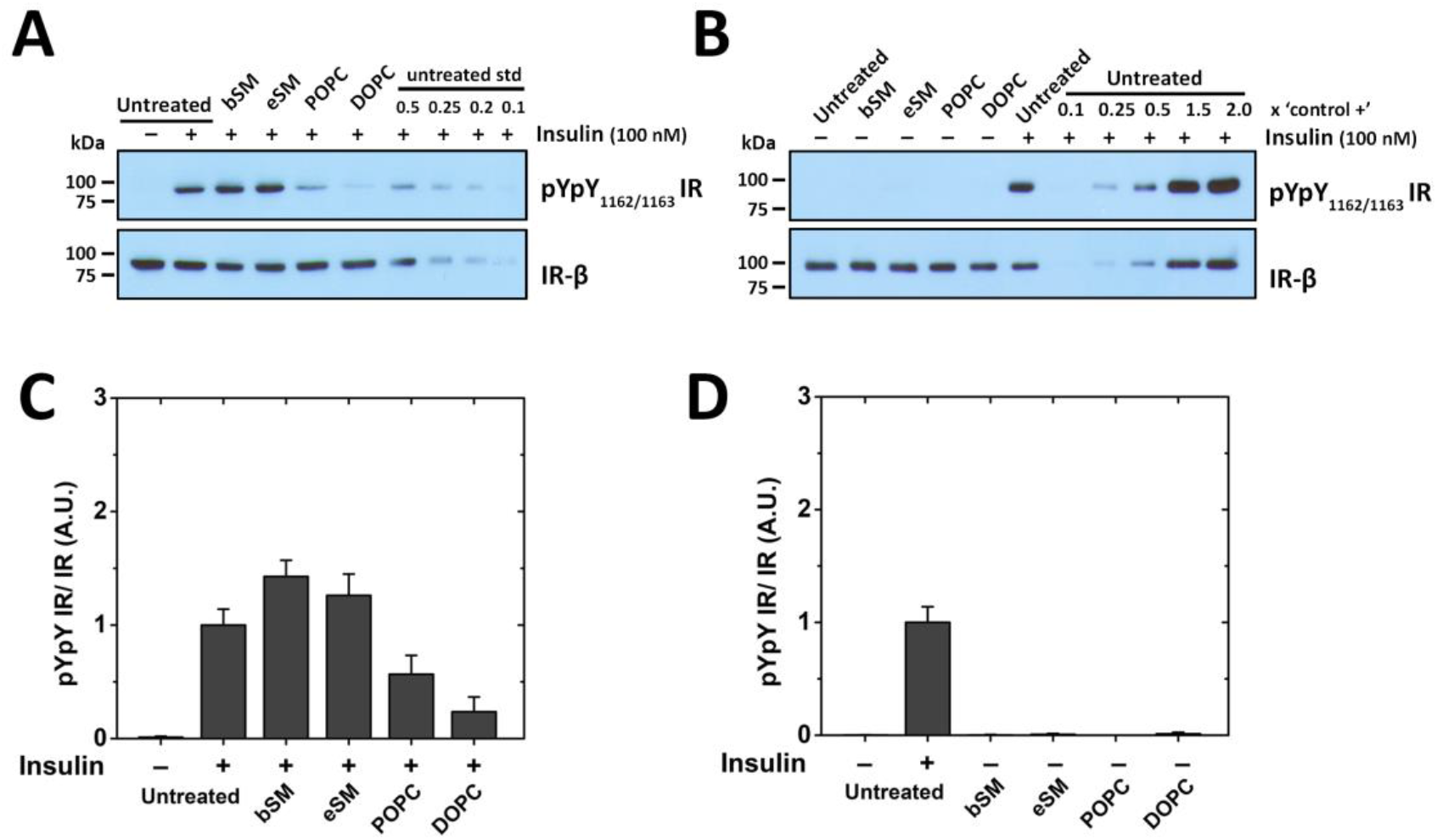
IR autophosphorylation activity after lipid exchange on CHO IR cells with SMs and unsaturated PCs. A representative western blot of (A) insulin stimulated cells and (B) unstimulated cells after lipid exchange with raft supporting bSM and eSM and non-raft supporting POPC and DOPC along with a standard curve of untreated insulin stimulated control is shown. A standard curve with different loadings of the untreated sample (from 0.1X to up to 2X) is included on the blots. Quantification of western blot data is shown for (C) stimulated and (D) unstimulated cells after lipid exchange. Band intensities of pYpY IR were normalized to IR-β band intensity for each sample. In this and the following experiments, the average (mean) and standard deviation from three independent experiments is shown, and activity in insulin stimulated untreated cells is defined as 1, unless otherwise noted.

Relative to untreated cells, insulin-stimulated samples show slightly higher IR autophosphorylation activity after exchange with brain or egg SM and reduced activity after exchange with unsaturated PCs **[Figure 2C]**. This reduction in IR autophosphorylation was most pronounced with DOPC, which is the lipid with the least ability to form ordered domains [35]–[38]. The correlation between IR activity and lipids that have a high propensity to form Lo domains is in agreement with our previous sterol substitution experiments [23]. No significant autophosphorylation activity was observed in unstimulated samples with either SMs or PCs **[Figure 2D]**.

### Changing plasma membrane bilayer width using lipid exchange shows higher IR activity in wider membranes

The lower IR autophosphorylation activity seen in Ld domain supporting lipids could be a consequence of narrow membrane width in an Ld bilayer compared to an Lo bilayer. An Lo bilayer forms wider bilayers than Ld because of close packing of lipids which minimizes gauche rotamers around C-C bonds [39]. To investigate the role of membrane width in more detail, we carried out lipid exchange with saturated PCs having varying acyl chain lengths: di12:0 PC (DLPC), di14 PC (DMPC), di16:0 PC (DPPC), and di18:0 PC (DSPC). Since we are only changing the outer leaflet lipids, we expect a change in membrane thickness of ∼ 4.8 Å from DLPC to DSPC, assuming lengthening an acyl chain leads to a 0.8 Å increase in bilayer width per carbon atom [40]. Estimated changes in plasma membrane widths for the Lo and Ld states after exchange with DLPC, DMPC, DPPC, and DSPC are shown in **Supplementary table 1**. These span a range from narrower to wider than on the unexchanged plasma membrane **[Supplementary table 1]**. In addition, experiments in asymmetric artificial lipid vesicles show that the formation of Lo domains increases monotonically as acyl chain length is increased, with DSPC containing the highest fraction of Lo domains [41].

**Figure 3A** shows that in insulin stimulated cells, IR autophosphorylation activity increases monotonically with acyl chain length. Compared to untreated cells, we observe a significantly lower IR activity with the shortest chain (DLPC), while the longest chain (DSPC) shows more than double the activity in untreated cells. After lipid exchange with saturated PCs, unstimulated cells show a low level of basal autophosphorylation higher than the unstimulated untreated cells **[Figure 3B]**. This suggests some loss of regulation of basal IR activity after introduction of saturated PCs.

**Figure 3:**
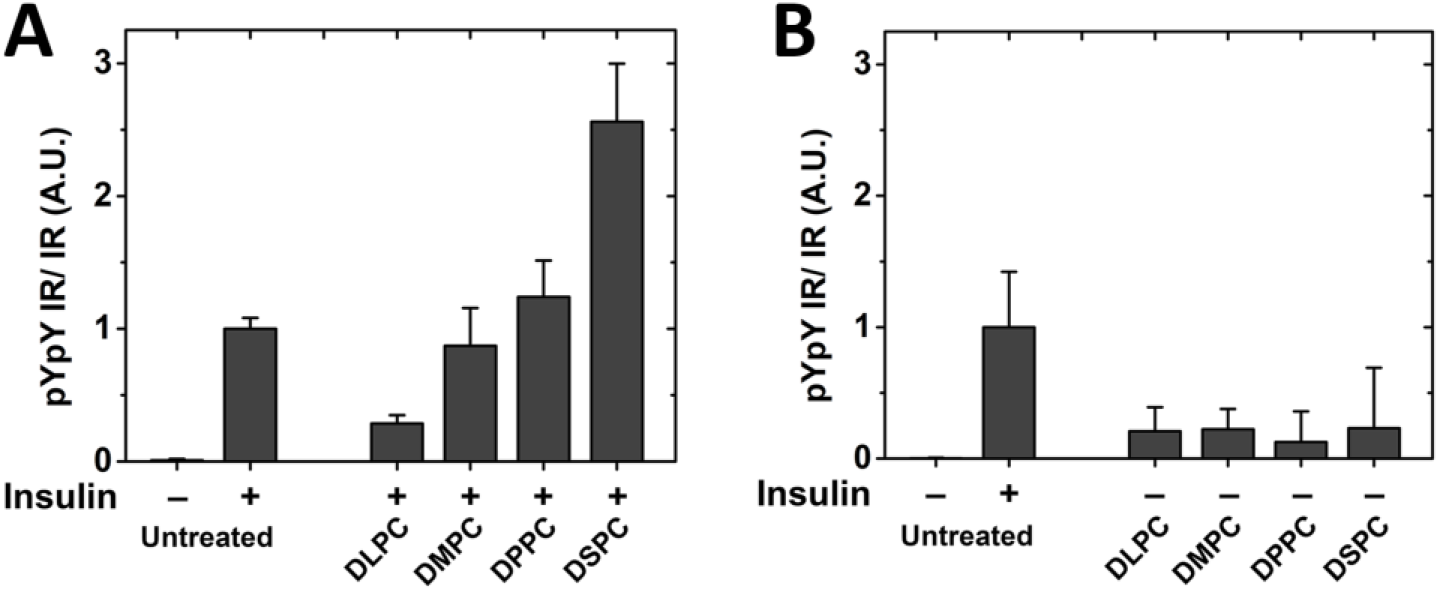
IR autophosphorylation activity after lipid exchange with saturated PCs of short and long chain lipids. Analysis of activity from western blot quantification of (A) stimulated and (B) unstimulated IR autophosphorylation activity after exchange with DLPC, DMPC, DPPC and DSPC along with untreated control.

### Strong correlation between propensity to form ordered state bilayers and increased IR activity

The observations above indicate that the effect of lipid exchange on the propensity of lipids to form the Lo state and its effect on bilayer width both regulate IR activity. Since saturated PCs that form wider bilayers have an increased propensity to form an ordered state, it is possible that the effect of lipid substitutions largely reflect degree to which lipid substitution enhances the ability to form the Lo state. A strong correlation between the ability to form an ordered state and IR activity supports this hypothesis. This correlation can be seen clearly in **Figure 4** in which IR autophosphorylation activity is graphed against the gel-to-Ld phase transition temperatures (Tm) **[Supplementary table 2]** of the lipids in vesicles containing a single lipid species. The gel state is highly ordered, and in artifical asymmetric membranes high Tm is associated with a greater tendency to form Lo domains when cholesterol is present [42].

**Figure 4:**
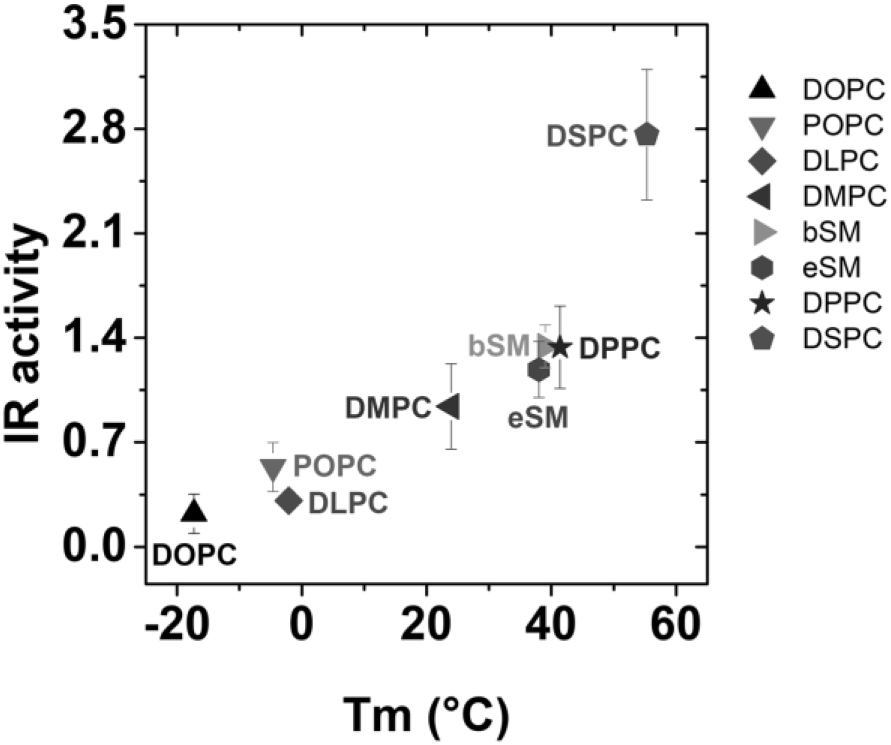
Insulin stimulated IR autophosphorylation activity compiled from **Figures 2** and **3** vs. gel to Ld state lipid phase transition temperatures of lipid vesicles composed of pure exchanged lipid. Activity in untreated cells has been normalized to a value of 1.

### IR activity in alkyl maltoside micelles of increasing micelle width shows higher IR activity in wider micelles

To test if there is also a direct effect of membrane width on IR activity, we measured the dependence of IR activity upon hydrophobic width under conditions in which there is no domain formation. This was done by incorporating purified full-length IR in alkyl maltoside micelles with increasing alkyl chain lengths (from 10 to 16 carbons). The detergents used were n-decyl-β-D maltopyranoside (DM), n-dodecyl-β-D maltopyranoside (DDM), n-tetradecyl-β-D maltopyranoside (TDM), and n-hexadecyl-β-D-maltopyranoside (HDM). Alkyl maltoside detergents are reported to form oblate ellipsoid structures with measured (or estimated) minor axis dimensions (widths) of 27 Å, 32 Å, 37 Å and 41 Å for DM, DDM, TDM and HDM respectively [43] **[Supplementary table 3]**. For reference, the plasma membrane Ld and Lo thicknesses are approximately 33 Å and 37 Å, respectively [40], and for a homogeneous plasma membrane bilayer in which the outer leaflet is fully substituted with DLPC, DMPC, DPPC or DSPC, we estimate bilayer widths of 30 to 39 Å **[Supplementary table 1]** depending on the physical state of the membrane. Thus, the detergent micelles roughly span the hydrophobic widths examined in saturated PC exchange experiments.

To carry out the experiments in detergent, IR was first affinity purified in DDM lysis buffer at 0.29 mM (critical micelle concentration (CMC) of DDM is 0.17 mM). The IR dissolved in DDM was diluted 1:10 with each of the four detergents, which were at concentrations of (0.5 + CMC) mM, to ensure 0.5 mM of detergent would be in the form of micelles in all cases **[Supplementary table 3]**. The DDM present after dilution should not significantly alter the average alkyl chain length. Even if all the diluted DDM is incorporated into the micelles of DM, TDM, and HDM, it would alter hydrophobic width by less than 0.5 Å.

Insulin-stimulated IR autophosphorylation activity measured by western blotting with an anti-phosphotyrosine antibody shows an increase with increasing micelle width **[Figure 5A]**. The difference between the narrowest and widest micelles is about 2-fold. This is much less than the change in activity observed when membrane width was altered by lipid exchange. For wider micelles, the activity increase is also seen in samples lacking insulin stimulation **[Figure 5A]**.

**Figure 5:**
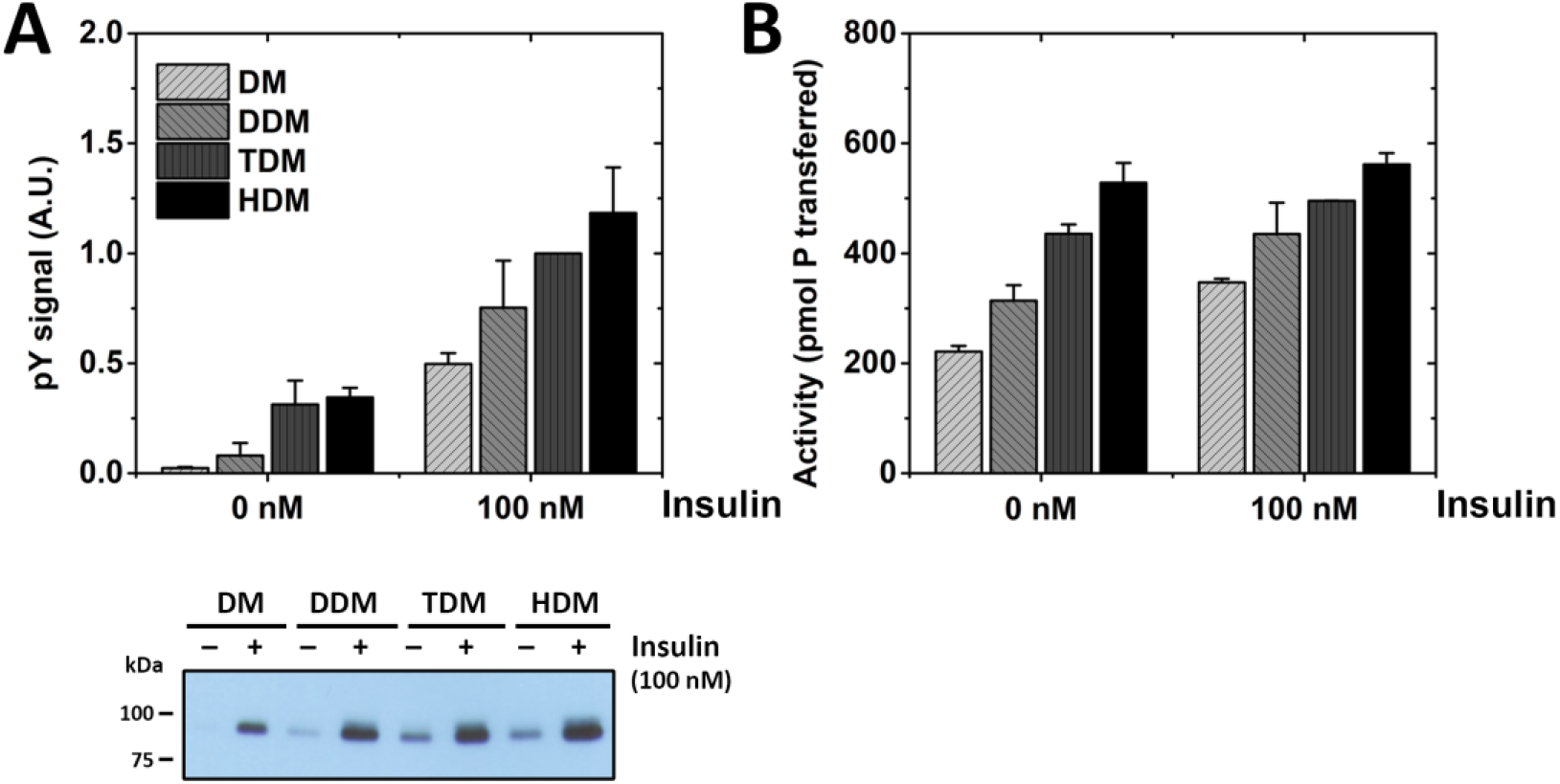
IR kinase activity increases in maltoside micelles of increasing micelle width. (A) IR autophosphorylation activity measured in maltoside micelles by western blot using phosphotyrosine antibody with and without 100 nM insulin. Densitometry analysis of bands from three independent experiments normalized to ‘100 nM TDM’ in each blot is shown. A representative sample blot is depicted below. (B) Radioactive kinase assay of IR activity on peptide substrate E_4_YM_4_ with and without 100 nM insulin stimulation measured as pico-mol phosphate transferred to peptide from three independent experiments is shown.

We also assayed ATP-dependent IR-catalyzed phosphorylation of the synthetic peptide substrate E_4_YM_4_ (KKEEEEYMMMMG) in the alkyl maltoside micelles **[Figure 5B]**. The activity of the receptor toward peptide substrate showed the same trend of increasing activity with increasing micelle width, but the difference between the thinnest and widest micelles was only 1.5-fold. In this assay, activity was only slightly higher in the presence of insulin relative to that in its absence. Thus, both the autophosphorylation and peptide phosphorylation data indicate that the formation of a substantial level of the active IR conformation does not require insulin binding in detergent micelles.

### Global inhibition of phosphotyrosine phosphatases show phosphatases are partially responsible for the change in IR activity observed after lipid exchanges

The above experiments indicate that IR is considerably activated when membranes have a greater propensity to form Lo domains, but that an increase in membrane width does not fully explain the large increase in IR activity. We next examined if a lipid-dependent decrease in accessibiiity to phosphatases was involved. Phosphorylation levels in cells are a dynamic interplay between kinases and phosphatases [44], [45]. For other receptor kinases believed to be activated by localization in Lo domains, segregation from phosphatases located in Ld domains is thought to be an important factor in enhancement of kinase activity [46], [47]. If IR localization in Lo domains has a similar effect on accessibility to phosphatases, it would be predicted that phosphatases have less effect when the lipid substitutions increase the propensity to form Lo domains.

To examine the effect of phosphatases on autophosphorylation, we compared activity after lipid exchange experiments in the absence and presence of a global phosphotyrosine phosphatase inhibitor, sodium orthovanadate (SOV) [30]. **Figure 6A** shows a partial rescue of insulin-stimulated IR autophosphorylation activity after POPC and DOPC exchange in the presence of SOV. However, the IR activity is not fully restored in POPC and DOPC when compared to the SOV treated control (untreated by lipid exchange) and to the SM exchanged samples. It is not clear whether this reflects lower intrinsic kinase activity or incomplete inhibition of phosphatases. Unstimulated samples showed no significant activity after lipid exchange when SOV was present **[Figure 6B]**.

**Figure 6:**
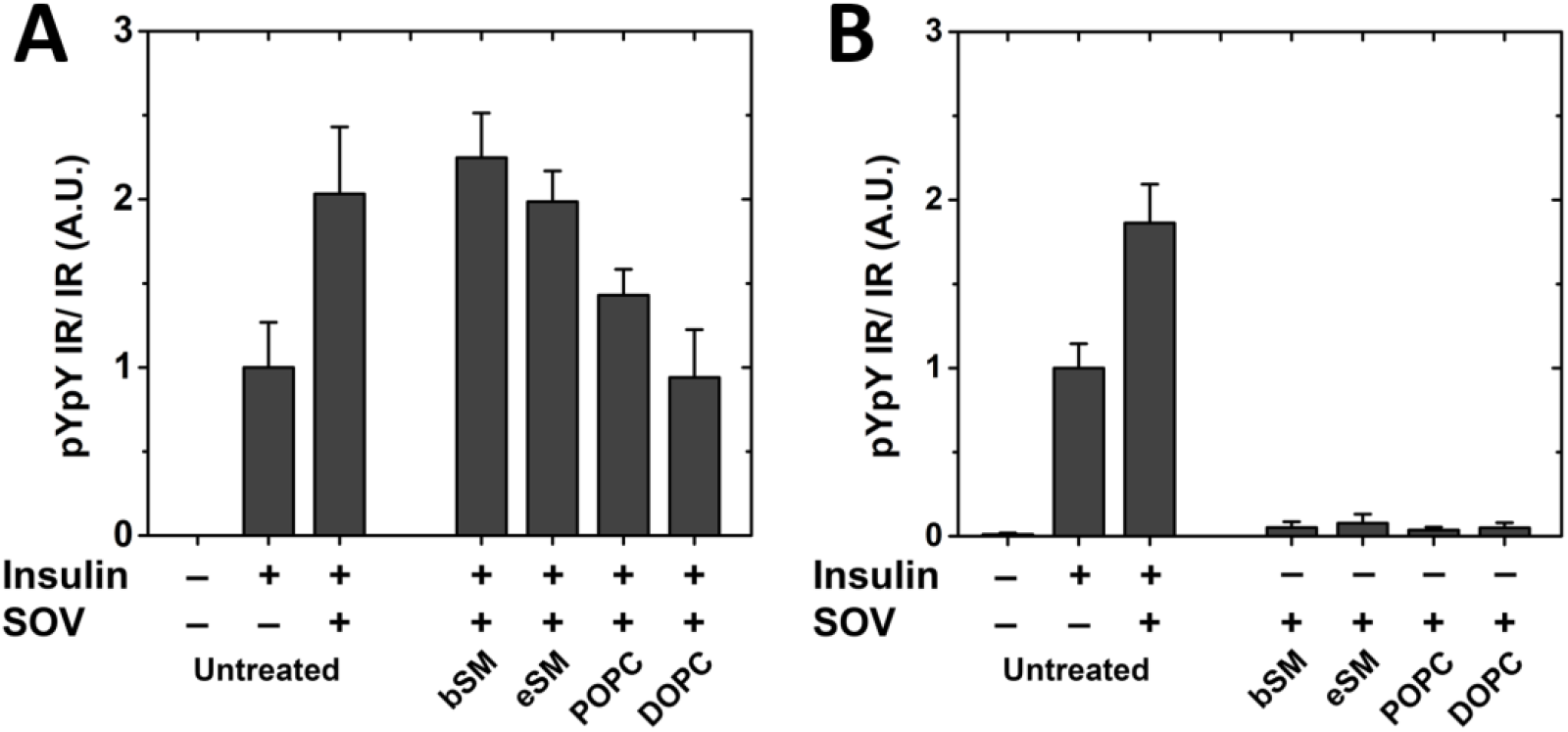
IR activity in the presence of SOV after lipid exchange using SMs and unsaturated PCs. Activity of insulin-stimulated (A) and unstimulated (B) cells after lipid exchange is shown.

There was also an effect of global phosphatase inhibition using SOV on cells that had undergone substitution with various saturated PCs. **Figure 7A** shows that after insulin stimulation in SOV-treated cells, the effect of acyl chain length upon activity was much smaller than in the absence of SOV **(see Figure 3**).

**Figure 7:**
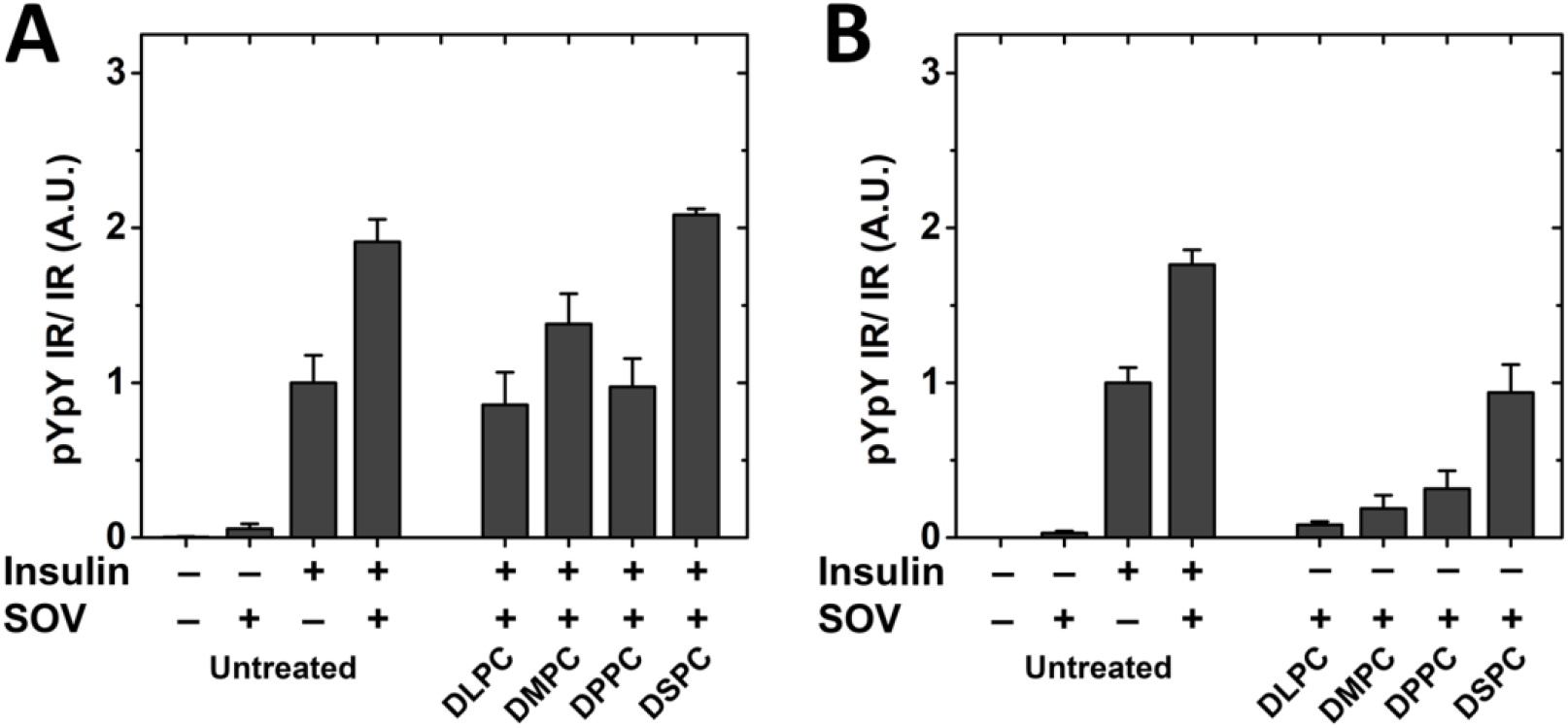
IR activity in the presence of SOV after lipid exchange with saturated PCs of different chain lengths. Activity of insulin-stimulated (A) and unstimulated cells (B) after treatment with SOV is shown.

Interestingly, in the presence of SOV, there was an increase in unstimulated IR activity with increasing PC acyl chain length **[Figure 7B]**. DSPC exchanged cells had an activity similar to that in unexchanged insulin-stimulated cells in the absence of insulin. This shows that a very wide membrane is sufficient to activate IR in the absence of insulin.

To better illustrate the effect of phosphatases on IR activity, we graphed the relative effect of phosphatases (IR activity without SOV divided by IR activity with SOV) vs Tm **[Figure 8]**. A value of 1 indicates no significant difference in IR activity in the presence (-SOV) or absence of phosphatases (+SOV), while a value below 1 suggests a higher IR activity in the absence of phosphatases (+SOV) indicating phosphatases have a substantial role in removing the activity of IR after exchange. Low Tm lipids such as DOPC, DLPC and POPC that have a lower propensity to form Lo domains have values below 0.5 showing the greatest effect after phosphatase inhibition **[Figure 8]**. This might indicate a higher accessibility of phosphatases to IR in Ld domains.

**Figure 8:**
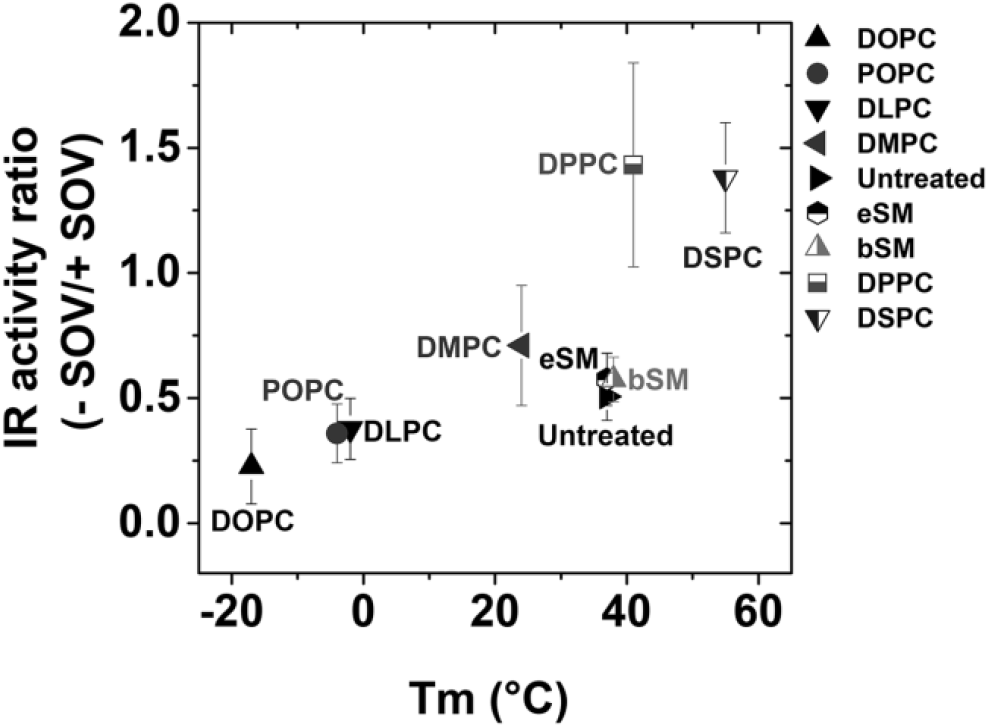
Correlation of relative effect of phosphatases on IR activity with gel-to-Ld Tm phase transition temperature of exchanged lipid. IR autophosphorylation activity without SOV is divided by IR activity with SOV treatment to illustrate the relative effect of phosphatases. A low value indicates a high degree of accessibility to phosphatases.

Since IR activity was measured at room temperature, about 23°C, it is possible that the observed effect of lipid domains on IR activity was more pronounced than at physiological temperature because lipid Lo domain formation is favored at lower temperatures [48]. To test this, activity and subsequent processing was measured at 37°C. We carried out these experiments on untreated cells and using as representative lipids DOPC (lipid with least propensity to form Lo domains) and DSPC (lipid with the highest propensity to form Lo domains). Similar to the results shown in **Figure 2** and **Figure 3**, IR autophosphorylation activity was low after DOPC exchange and high after DSPC exchange, compared to the untreated control **[Supplementary figure 4]**. Treatment with SOV shows that at room temperature and at 37 °C the largest increase in activity was measured in DOPC-exchanged samples, intermediate increase in activity in untreated samples and no increase in activity in DSPC-exchanged samples **[Supplementary figure 4]**. These results suggest that the observed IR activity responses to lipid exchanges are relevant at physiological temperatures.

## DISCUSSION

Our previous study using sterol replacement after cholesterol depletion suggested that sterols supporting ordered domains provide a favorable environment for IR to be activated [23]. In this study, the role of lipid organization on IR autophosphorylation activity in cells was investigated by manipulating the plasma membrane outer leaflet using MαCD mediated lipid exchange [26], [27]. In agreement with the sterol results, insulin-stimulated IR autophosphorylation activity is higher in plasma membranes exchanged with lipids that have a higher propensity to form wide bilayers and Lo domains (high Tm lipids). We also observed a direct membrane thickness effect on IR activity, with wider maltoside micelle hydrophobic widths supporting higher activity of purified IR. Combined with the fact that Lo domains are wider than Ld domains, these experiments are consistent with the model that increased association of IR with Lo domains is responsible for the increased IR activity in lipids that tend to support Lo domains.

An alternative model is that before and after exchange the PM remains homogeneous, lacking domains, and that the effect of activating IR after lipid exchange is due to an increase in bilayer width. This alternative model is less likely to be correct for several reasons. First, the activating effect of increasing PC hydrocarbon chain length is much more than the effect of changing detergent hydrocarbon chain length. If there are no domains, one would predict they should have had similar levels of activation. Second, studies in plasma membrane vesicles formed after lipid exchange and in artificial asymmetric lipid vesicles show that lipid compositions increasing the propensity to form Lo domains do result in increased formation of ordered lipid domains under physiologic or near physiologic conditions [42], [48]. Third, the domain model is favored by the close relationship between decreased sensitivity of IR activity to phosphatases in lipid substitutions that favor Lo domain formation. This suggests a sequestration where IR in Lo domains is protected from phosphatases **[Figure 9]**. It has been shown in other signal transduction systems that activation of receptor complexes with kinase activity is enhanced by segregation of phosphatases in Ld domains and kinases in Lo domains. For example, activated Lyn kinase is sequestered in ordered Lo domains, where it is protected from the action of phosphatases, which reside in Ld domains [49]. Additionally, or alternatively, IR in an Lo state bilayer may exist in a conformation that is inaccessible to phosphatases.

**Figure 9:**
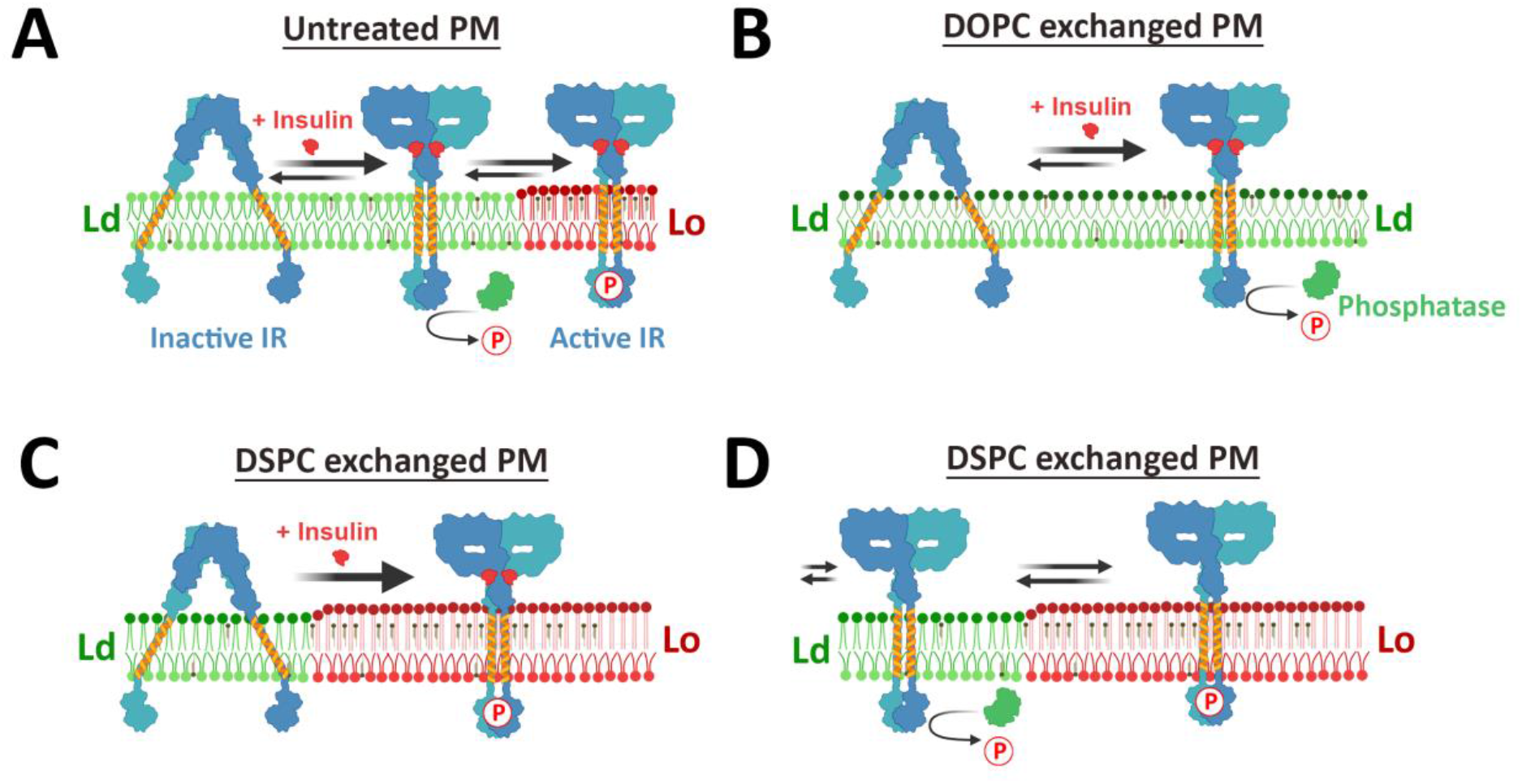
Schematic representation of the hypothesis for the effect of altering lipid composition upon IR activity. Green represents Ld domains, red Lo domains. Phosphate is shown as P. Note: inactive IR is shown with tilted TM domains, while the active has an untilted conformation. (A) In normal (untreated) PM, insulin binding promotes a conformational change that brings the kinase domains close together. This active conformation is supported in Lo domains where IR is protected from phosphatases and autophosphorylation proceeds. In Ld domains, phosphatases suppress the IR autophosphorylation. (B) In DOPC exchanged membranes with largely Ld environment, IR is not protected from phosphatases, resulting in reduced autophosphorylation. (C) In DSPC-exchanged membranes which are largely Lo domain forming with wider membranes, the untilted active conformation of insulin stimulated IR is favored and strongly localized in Lo domains and therefore protected from phosphatase activity. (D) Even in the absence of insulin in DSPC-exchanged membranes, IR can form its active conformation, and dynamically move between the Lo and Ld domains, with IR inactivated by phosphatases when it moves to the Ld domain.

In this regard, the behavior of IR after substitution with DSPC is especially noteworthy. After DSPC substitution, which should increase bilayer width to a value larger than in unexchanged cells **[Supplemental Table 1]**, the activity of IR is higher than in unexchanged cells **[Figure 3A]**. Furthermore, there is no effect of inhibiting phosphatases on insulin-stimulated IR activity after DSPC exchange. This is consistent with a model in which activated IR is totally sequestered in Lo domains after DSPC substitution **[Figure 9D]**. Even in the absence of insulin stimulation, IR is very active after DSPC exchange when phosphatases are inhibited **[Figure 7B]**. This suggests a scenario where unstimulated IR dynamically shifts between Lo and Ld domains, and IR phosphorylation is removed by phosphatases when it is in Ld domains. Insulin addition likely helps to lock the active conformation of IR and thus allows for IR to be in Lo domains longer for signal transduction, reducing the dynamic shift.

This activation of IR in wider membranes could be a consequence of a change in IR transmembrane (TM) helix orientation within the membrane in the active form vs inactive form. IR in its inactivated conformation takes on an ‘inverted V-shape’ form [50], which would be tilted relative to the plane of the membrane [**Figure 9**], and therefore might favor formation of a tilted TM helix [50]. The active ‘T-shaped dimer’ conformation which would be untilted with respect to the plane of the membrane, would be more likely to favor a TM helix that is not tilted. The TM helix of IR is about 23 residues long, so it could span the hydrophobic core of a bilayer of 34.5 Å without tilting. In narrower membranes, one would expect IR TM helices to be tilted to fit the hydrophobic width of the bilayer. In wider membranes such as DSPC (estimated 38.6 Å in the Lo state) the untilted TM helix would be strongly favored (perhaps with some accompanying local narrowing of the bilayer) [51]. Thus, in a wider bilayer the untilted helices could tend to favor the T-shaped dimer, especially if the connection between extracellular and TM domains is very rigid. This could also explain activating IR even in the absence of insulin after DSPC exchange, which would result in the widest bilayer width. Consistent with this model, it has been shown in previous studies that long TM segment length can favor a stronger affinity of TM proteins for Lo domains [52], [53], and this is due to the wider bilayer in the Lo domains [52]. This model is illustrated in **Figure 9**.

Thus, the results of lipid exchange studies in cells strongly favor a model of IR activity regulation where a combination of increased bilayer width and a relative inaccessibility of phosphatases in Lo domains favor IR activation. It is possible that alterations of lipid by diet and cell changes related to differentiation [54] might be sufficient to alter IR activity in physiological settings. This model for kinase regulation may also be applicable to other signal transduction processes in which kinases have a single TM helix.

## Notes

This work was supported by NIH grant GM 122493 (to E.L.) and Dept. of Veterans Affairs Merit Review Grant BX002292 (to W.T.M.)

### Competing Interest Statement

The authors have declared no competing interest.

## REFERENCES

[1] I. Levental, K. R. Levental, and F. A. Heberle, “Lipid Rafts: Controversies Resolved, Mysteries Remain,” Trends Cell Biol., vol. 30, no. 5, pp. 341–353, 2020, doi: 10.1016/j.tcb.2020.01.009.

[2] E. London, “How principles of domain formation in model membranes may explain ambiguities concerning lipid raft formation in cells,” Biochim. Biophys. Acta - Mol. Cell Res., vol. 1746, no. 3, pp. 203–220, 2005, doi: 10.1016/j.bbamcr.2005.09.002.

[3] D. Lingwood and K. Simons, “Lipid rafts as a membrane-organizing principle,” Science (80-.)., vol. 327, no. 5961, pp. 46–50, 2010, doi: 10.1126/science.1174621.

[4] L. J. Pike, “Growth factor receptors, lipid rafts and caveolae: An evolving story,” Biochim. Biophys. Acta - Mol. Cell Res., vol. 1746, no. 3, pp. 260–273, 2005, doi: 10.1016/j.bbamcr.2005.05.005.

[5] P. Varshney, V. Yadav, and N. Saini, “Lipid rafts in immune signalling: current progress and future perspective,” Immunology, vol. 149, no. 1, pp. 13–24, 2016, doi: 10.1111/imm.12617.

[6] D. A. Brown and E. London, “Functions of Lipid Rafts in Biological Membranes,” Annu. Rev. Cell Dev. Biol., vol. 14, no. 1, pp. 111–136, 1998, doi: 10.1146/annurev.cellbio.14.1.111.

[7] M. B. Stone, S. A. Shelby, M. F. Nńñez, K. Wisser, and S. L. Veatch, “Protein sorting by lipid phase-like domains supports emergent signaling function in b lymphocyte plasma membranes,” Elife, vol. 6, pp. 1–33, 2017, doi: 10.7554/eLife.19891.

[8] C. Y. Lin, J. Y. Huang, and L. Lo, “Unraveling the impact of lipid domains on the dimerization processes of single-molecule EGFRs of live cells,” BBA - Biomembr., vol. 1848, no. 3, pp. 886– 893, 2015, doi: 10.1016/j.bbamem.2014.12.019.

[9] R. A. Haeusler, T. E. Mcgraw, and D. Accili, “Biochemical and cellular properties of insulin receptor signalling,” Nat. Publ. Gr., vol. 19, no. 1, pp. 31–44, 2017, doi: 10.1038/nrm.2017.89.

[10] J. Boucher, A. Kleinridders, and C. Ronald Kahn, “Insulin receptor signaling in normal and insulin-resistant states,” Cold Spring Harb. Perspect. Biol., vol. 6, no. 1, pp. 1–23, 2014, doi: 10.1101/cshperspect.a009191.

[11] P. E. Bickel, “Lipid rafts and insulin signaling,” Am. J. Physiol. - Endocrinol. Metab., vol. 282, no. 1 45–1, 2002, doi: 10.1152/ajpendo.2002.282.1.e1.

[12] M. Karlsson et al., “Colocalization of insulin receptor and insulin receptor substrate-1 to caveolae in primary human adipocytes: Cholesterol depletion blocks insulin signalling for metabolic and mitogenic control,” Eur. J. Biochem., vol. 271, no. 12, pp. 2471–2479, 2004, doi: 10.1111/j.1432-1033.2004.04177.x.

[13] S. Parpal, M. Karlsson, H. Thorn, and P. Strålfors, “Cholesterol Depletion Disrupts Caveolae and Insulin Receptor Signaling for Metabolic Control via Insulin Receptor Substrate-1, but Not for Mitogen-activated Protein Kinase Control,” J. Biol. Chem., vol. 276, no. 13, pp. 9670–9678, 2001, doi: 10.1074/jbc.M007454200.

[14] J. Sánchez-Wandelmer et al., “Inhibition of cholesterol biosynthesis disrupts lipid raft/caveolae and affects insulin receptor activation in 3T3-L1 preadipocytes,” Biochim. Biophys. Acta - Biomembr., vol. 1788, no. 9, pp. 1731–1739, 2009, doi: 10.1016/j.bbamem.2009.05.002.

[15] N. Khatibzadeh, A. A. Spector, W. E. Brownell, and B. Anvari, “Effects of Plasma Membrane Cholesterol Level and Cytoskeleton F-Actin on Cell Protrusion Mechanics,” PLoS One, vol. 8, no. 2, 2013, doi: 10.1371/journal.pone.0057147.

[16] L. Zhang, L. Zhao, P. K. Ouyang, and P. Chen, “Insight into the role of cholesterol in modulation of morphology and mechanical properties of CHO-K1 cells: An in situ AFM study,” Front. Chem. Sci. Eng., vol. 13, no. 1, pp. 98–107, 2019, doi: 10.1007/s11705-018-1775-y.

[17] B. Hissa et al., “Membrane cholesterol removal changes mechanical properties of cells and induces secretion of a specific pool of lysosomes,” PLoS One, vol. 8, no. 12, 2013, doi: 10.1371/journal.pone.0082988.

[18] S. Vainio et al., “Dynamic association of human insulin receptor with lipid rafts in cells lacking caveolae.,” EMBO Rep., vol. 3, no. 1, pp. 95–100, 2002, doi: 10.1093/embo-reports/kvf010.

[19] J. Gustavsson et al., “Localization of the insulin receptor in caveolae of adipocyte plasma membrane,” FASEB J., vol. 13, no. 14, pp. 1961–1971, 1999, doi: 10.1096/fasebj.13.14.1961.

[20] D. A. Brown, “Lipid rafts, detergent-resistant membranes, and raft targeting signals,” Physiology, vol. 21, no. 6, pp. 430–439, 2006, doi: 10.1152/physiol.00032.2006.

[21] D. Lichtenberg, F. M. Goñi, and H. Heerklotz, “Detergent-resistant membranes should not be identified with membrane rafts,” Trends Biochem. Sci., vol. 30, no. 8, pp. 430–436, 2005, doi: 10.1016/j.tibs.2005.06.004.

[22] J. H. Kim and E. London, “Using Sterol Substitution to Probe the Role of Membrane Domains in Membrane Functions,” Lipids, vol. 50, no. 8, pp. 721–734, 2015, doi: 10.1007/s11745-015-4007-y.

[23] R. J. Delle Bovi, J. H. Kim, P. Suresh, E. London, and W. T. Miller, “Sterol structure dependence of insulin receptor and insulin-like growth factor 1 receptor activation,” Biochim. Biophys. Acta - Biomembr., vol. 1861, no. 4, pp. 819–826, 2019, doi: 10.1016/j.bbamem.2019.01.009.

[24] M. Amoui, B. P. Craddock, and W. T. Miller, “Differential phosphorylation of IRS-1 by insulin and insulin-like growth factor I receptors in Chinese hamster ovary cells,” pp. 153–162, 2001.

[25] R. J. Delle Bovi and W. T. Miller, “Expression and purification of functional insulin and insulin-like growth factor 1 holoreceptors from mammalian cells,” Anal. Biochem., vol. 536, pp. 69–77, 2017, doi: 10.1016/j.ab.2017.08.011.

[26] G. Li, J. Kim, Z. Huang, J. R. St. Clair, D. A. Brown, and E. London, “Efficient replacement of plasma membrane outer leaflet phospholipids and sphingolipids in cells with exogenous lipids,” Proc. Natl. Acad. Sci., vol. 113, no. 49, pp. 14025–14030, 2016, doi: 10.1073/pnas.1610705113.

[27] G. Li, S. Kakuda, P. Suresh, D. Canals, S. Salamone, and E. London, “Replacing plasma membrane outer leaflet lipids with exogenous lipid without damaging membrane integrity,” PLoS One, vol. 14, no. 10, pp. 1–22, 2019, doi: 10.1371/journal.pone.0223572.

[28] U. K. Laemmli, “Cleavage of Structural Proteins during the Assembly of the Head of Bacteriophage T4,” vol. 227, pp. 680–685, 1970.

[29] M. M. Bradford, “A Rapid and Sensitive Method for the Quantitation Microgram Quantities of Protein Utilizing the Principle of Protein-Dye Binding,” vol. 254, pp. 248–254, 1976.

[30] J. A. Gordon, “Use of Vanadate as Protein - Phosphotyrosine Phosphatase Inhibitor,” vol. 20, no. 1982, 1991, pp. 477–482.

[31] C. B. Baron and R. F. Coburn, “Comparison of two copper reagents for detection of saturated and unsaturated neutral lipids by charring densitometry,” J. Liq. Chromatogr., vol. 7, no. 14, pp. 2793– 2801, 1984, doi: 10.1080/01483918408067046.

[32] Q. Lin and E. London, “Preparation of artificial plasma membrane mimicking vesicles with lipid asymmetry,” PLoS One, vol. 9, no. 1, 2014, doi: 10.1371/journal.pone.0087903.

[33] J. L. Symons et al., “Lipidomic atlas of mammalian cell membranes reveals hierarchical variation induced by culture conditions, subcellular membranes, and cell lineages,” Soft Matter, vol. 17, no. 2, pp. 288–297, 2021, doi: 10.1039/d0sm00404a.

[34] H. T. Cheng, Megha, and E. London, “Preparation and properties of asymmetric vesicles that mimic cell membranes. Effect upon lipid raft formation and transmembrane helix orientation,” J. Biol. Chem., vol. 284, no. 10, pp. 6079–6092, 2009, doi: 10.1074/jbc.M806077200.

[35] D. A. Brown and E. London, “Structure and origin of ordered lipid domains in biological membranes,” J. Membr. Biol., vol. 164, no. 2, pp. 103–114, 1998, doi: 10.1007/s002329900397.

[36] M. Mihailescu et al., “Acyl-chain methyl distributions of liquid-ordered and -disordered membranes,” Biophys. J., vol. 100, no. 6, pp. 1455–1462, 2011, doi: 10.1016/j.bpj.2011.01.035.

[37] R. Koynova and M. Caffrey, “Phases and phase transitions of the phosphatidylcholines,” Biochim. Biophys. Acta - Rev. Biomembr., vol. 1376, no. 1, pp. 91–145, 1998, doi: 10.1016/S0304-4157(98)00006-9.

[38] H. T. Cheng, Megha, and E. London, “Preparation and properties of asymmetric vesicles that mimic cell membranes. Effect upon lipid raft formation and transmembrane helix orientation,” J. Biol. Chem., vol. 284, no. 10, pp. 6079–6092, 2009, doi: 10.1074/jbc.M806077200.

[39] F. A. Heberle et al., “Bilayer thickness mismatch controls domain size in model membranes,” J. Am. Chem. Soc., vol. 135, no. 18, pp. 6853–6859, 2013, doi: 10.1021/ja3113615.

[40] F. A. Heberle, M. Doktorova, H. L. Scott, A. D. Skinkle, M. N. Waxham, and I. Levental, “Direct label-free imaging of nanodomains in biomimetic and biological membranes by cryogenic electron microscopy,” Proc. Natl. Acad. Sci. U. S. A., vol. 117, no. 33, pp. 19943–19952, 2020, doi: 10.1073/PNAS.2002200117.

[41] Q. Wang and E. London, “Lipid Structure and Composition Control Consequences of Interleaflet Coupling in Asymmetric Vesicles,” Biophys. J., vol. 115, no. 4, pp. 664–678, 2018, doi: 10.1016/j.bpj.2018.07.011.

[42] Q. Wang and E. London, “Lipid Structure and Composition Control Consequences of Interleaflet Coupling in Asymmetric Vesicles,” Biophys. J., vol. 115, no. 4, pp. 664–678, 2018, doi: 10.1016/j.bpj.2018.07.011.

[43] R. C. Oliver, J. Lipfert, D. A. Fox, R. H. Lo, S. Doniach, and L. Columbus, “Dependence of Micelle Size and Shape on Detergent Alkyl Chain Length and Head Group,” PLoS One, vol. 8, no. 5, 2013, doi: 10.1371/journal.pone.0062488.

[44] H. Sun and N. K. Tonks, “The coordinated action of protein tyrosine phosphatases and kinases in cell signaling,” Trends Biochem. Sci., vol. 19, no. 11, pp. 480–485, 1994, doi: 10.1016/0968-0004(94)90134-1.

[45] A. Cheng, N. Dubé, F. Gu, and M. L. Tremblay, “Coordinated action of protein tyrosine phosphatases in insulin signal transduction,” Eur. J. Biochem., vol. 269, no. 4, pp. 1050–1059, 2002, doi: 10.1046/j.0014-2956.2002.02756.x.

[46] R. M. Young, D. Holowka, and B. Baird, “A lipid raft environment enhances Lyn kinase activity by protecting the active site tyrosine from dephosphorylation,” J. Biol. Chem., vol. 278, no. 23, pp. 20746–20752, 2003, doi: 10.1074/jbc.M211402200.

[47] X. Su et al., “Phase separation of signaling molecules promotes T cell receptor signal transduction,” Science (80-.)., vol. 352, no. 6285, pp. 595–599, 2016, doi: 10.1126/science.aad9964.

[48] G. Li, Q. Wang, S. Kakuda, and E. London, “Nanodomains can persist at physiologic temperature in plasma membrane vesicles and be modulated by altering cell lipids,” J. Lipid Res., p. jlr.RA119000565, 2020, doi: 10.1194/jlr.ra119000565.

[49] R. M. Young, D. Holowka, and B. Baird, “A lipid raft environment enhances Lyn kinase activity by protecting the active site tyrosine from dephosphorylation,” J. Biol. Chem., vol. 278, no. 23, pp. 20746–20752, 2003, doi: 10.1074/jbc.M211402200.

[50] E. Uchikawa, E. Choi, G. Shang, H. Yu, and B. Xiao-Chen, “Activation mechanism of the insulin receptor revealed by cryo-EM structure of the fully liganded receptor-ligand complex,” Elife, vol. 8, pp. 1–23, 2019, doi: 10.7554/eLife.48630.

[51] S. S. Krishnakumar and E. London, “Effect of Sequence Hydrophobicity and Bilayer Width upon the Minimum Length Required for the Formation of Transmembrane Helices in Membranes,” J. Mol. Biol., vol. 374, no. 3, pp. 671–687, 2007, doi: 10.1016/j.jmb.2007.09.037.

[52] Q. Lin and E. London, “Altering hydrophobic sequence lengths shows that hydrophobic mismatch controls affinity for ordered lipid domains (rafts) in the multitransmembrane strand protein perfringolysin O,” J. Biol. Chem., vol. 288, no. 2, pp. 1340–1352, 2013, doi: 10.1074/jbc.M112.415596.

[53] B. B. Diaz-Rohrer, K. R. Levental, K. Simons, and I. Levental, “Membrane raft association is a determinant of plasma membrane localization,” Proc. Natl. Acad. Sci. U. S. A., vol. 111, no. 23, pp. 8500–8505, 2014, doi: 10.1073/pnas.1404582111.

[54] J. J. Lühr et al., “Maturation of Monocyte-Derived DCs Leads to Increased Cellular Stiffness, Higher Membrane Fluidity, and Changed Lipid Composition,” Front. Immunol., vol. 11, no. November, pp. 1–18, 2020, doi: 10.3389/fimmu.2020.590121.

[55] T. J. McIntosh, S. A. Simon, D. Needham, and C. hsien Huang, “Structure and Cohesive Properties of Sphingomyelin/Cholesterol Bilayers,” Biochemistry, vol. 31, no. 7, pp. 2012–2020, 1992, doi: 10.1021/bi00122a017.

[56] Z. Arsov, E. J. González-Ramírez, F. M. Goñi, S. Tristram-Nagle, and J. F. Nagle, “Phase behavior of palmitoyl and egg sphingomyelin,” Chem. Phys. Lipids, vol. 213, no. April, pp. 102– 110, 2018, doi: 10.1016/j.chemphyslip.2018.03.003.

[57] K. Tada, E. Miyazaki, M. Goto, N. Tamai, H. Matsuki, and S. Kaneshina, “Barotropic and thermotropic bilayer phase behavior of positional isomers of unsaturated mixed-chain phosphatidylcholines,” Biochim. Biophys. Acta - Biomembr., vol. 1788, no. 5, pp. 1056–1063, 2009, doi: 10.1016/j.bbamem.2009.02.008.

[58] R. N. Lewis, B. D. Sykes, and R. N. McElhaney, “Thermotropic Phase Behavior of Model Membranes Composed of Phosphatidylcholines Containing Cis-Monounsaturated Acyl Chain Homologues of Oleic Acid: Differential Scanning Calorimetric and31P NMR Spectroscopic Studies,” Biochemistry, vol. 27, no. 3, pp. 880–887, 1988, doi: 10.1021/bi00403a007.

[59] R. N. Lewis, N. M. Nanette Mak, and R. N. McElhaney, “A Differential Scanning Calorimetric Study of the Thermotropic Phase Behavior of Model Membranes Composed of Phosphatidylcholines Containing Linear Saturated Fatty Acyl Chains,” Biochemistry, vol. 26, no. 19, pp. 6118–6126, 1987, doi: 10.1021/bi00393a026.

