## Supplementary Figures and Tables for "Phospholipid exchange shows insulin receptor activity is supported by both the propensity to form wide bilayers and ordered domains (rafts)"

**List of materials included**: Supplementary figures 1, 2, 3, 4. Supplementary tables 1, 2, 3.

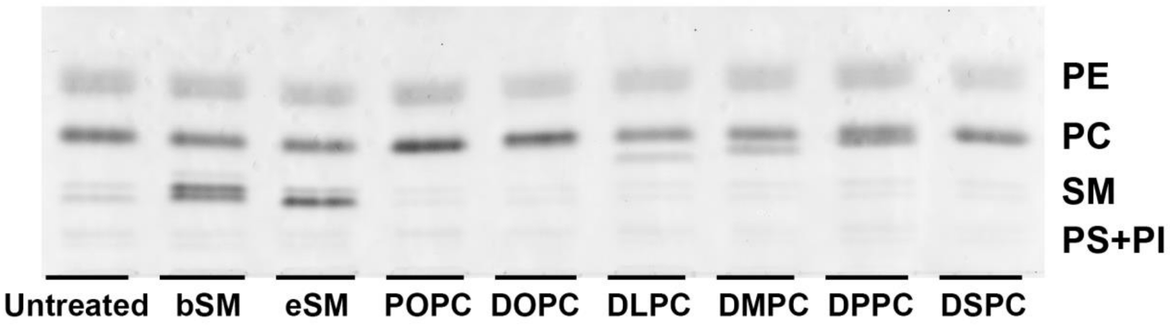


**Supplementary figure 1:** HP-TLC of lipid exchange in CHO IR cells. Representative HP-TLC image of untreated and lipid exchanged CHO IR cells using brain SM, egg SM, POPC, DOPC, DLPC DMPC, DPPC, and DSPC. Position of lipid bands corresponding to phosphatidylethanolamine (PE), phosphatidylcholine (PC), sphingomyelin (SM), phosphatidylserine (PS) and phosphatidylinositol (PI) are shown on the right. Note that the band for exogenous PC exchanged into cells can be detected under the main PC band for PC acyl chains are short (DLPC and DMPC.)

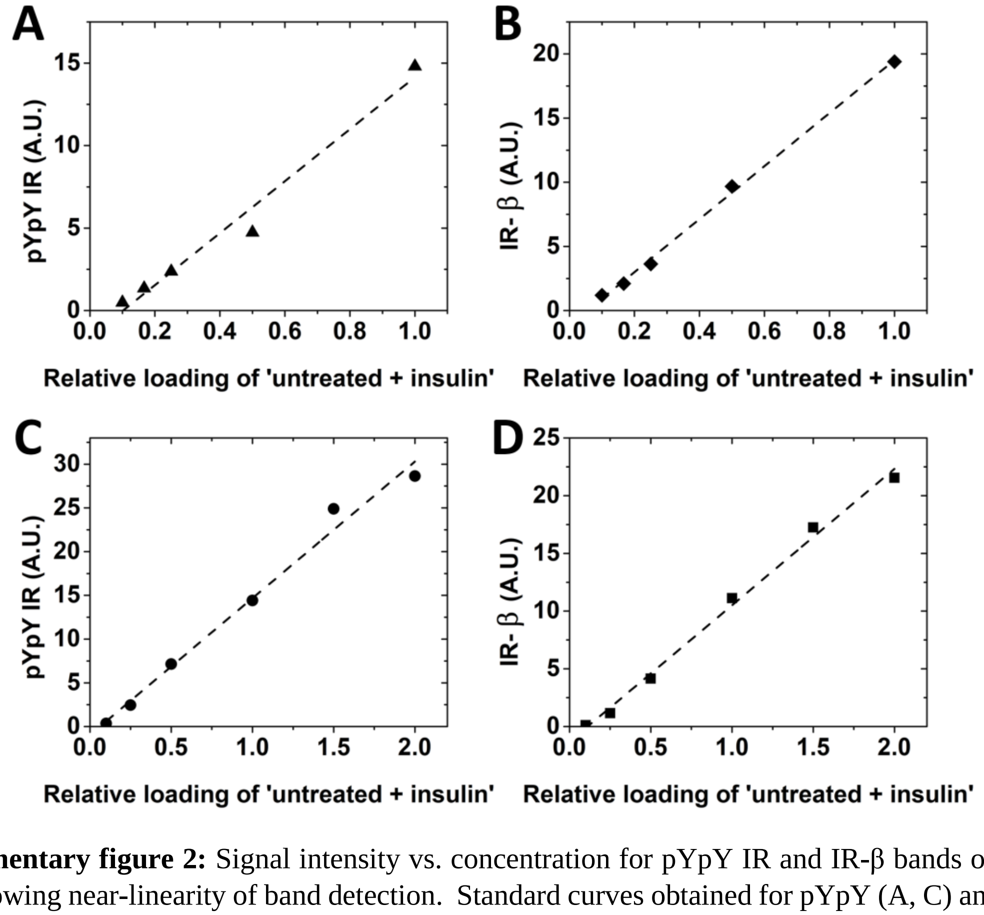


**Supplementary figure 2:** Signal intensity vs. concentration for pYpY IR and IR-β bands on Western blots showing near-linearity of band detection. Standard curves obtained for pYpY (A, C) and IR-β (B, D) bands for 'untreated + insulin' samples of the blots shown in **Figure 2A** and **2B** respectively.

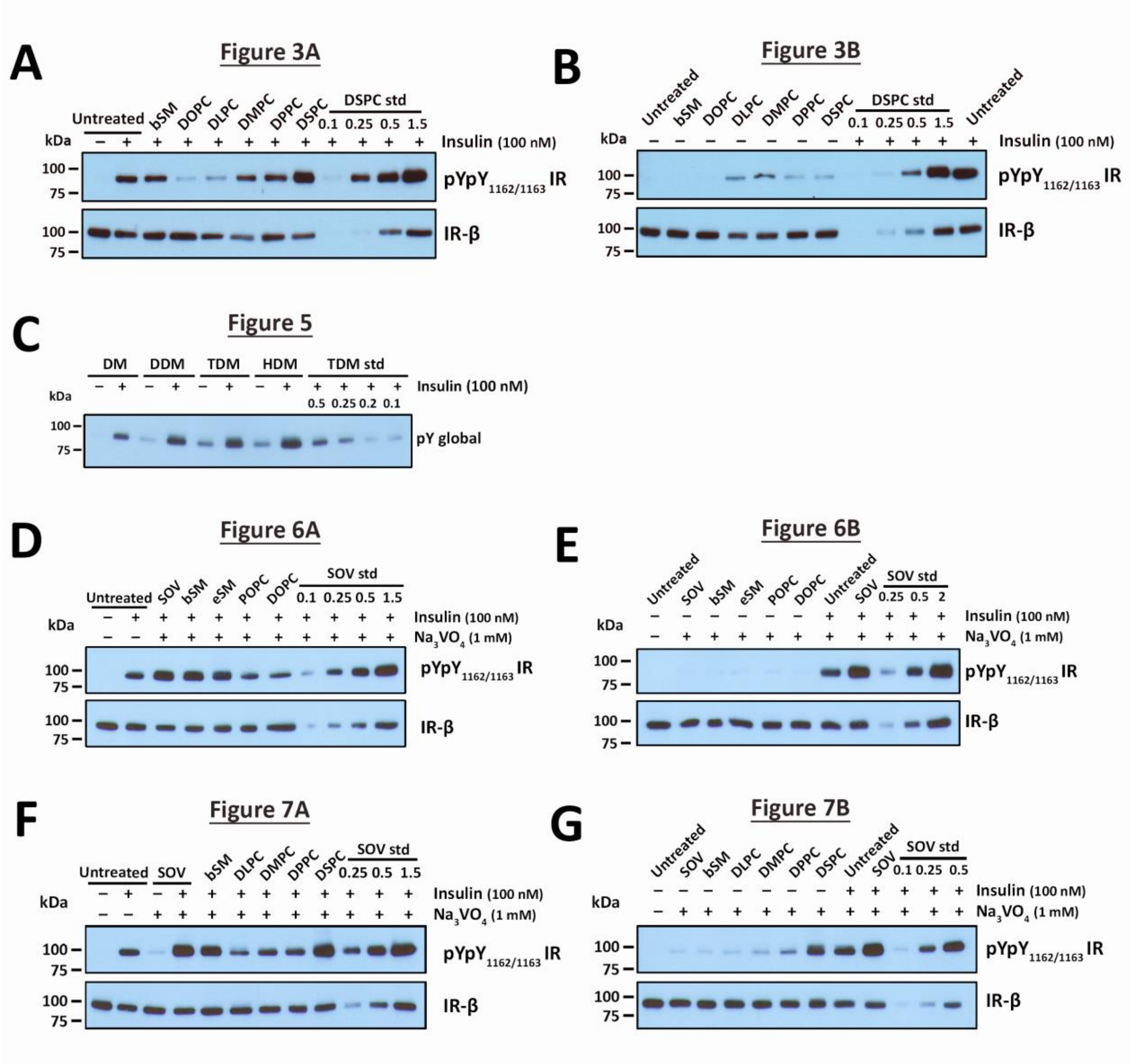


**Supplementary figure 3:** Representative western blot images after lipid exchange in CHO IR cells. Complete representative western blot images. Standard curves composed of either untreated control samples, or samples with highest intensity are shown in the rightmost lanes in each western blot image and labeled as 'std'. (A) **Figure 3** insulin-stimulated and (B) unstimulated lipid exchanges with saturated PCs with increasing acyl chain lengths. (C) **Figure 5** autophosphorylation of purified IR in alkyl maltoside micelles. (D) **Figure 6** stimulated and (E) unstimulated lipid exchange with SMs and PCs with SOV treatment. (F) **Figure 7** stimulated and (G) unstimulated lipid exchange using saturated PCs with SOV treatment.

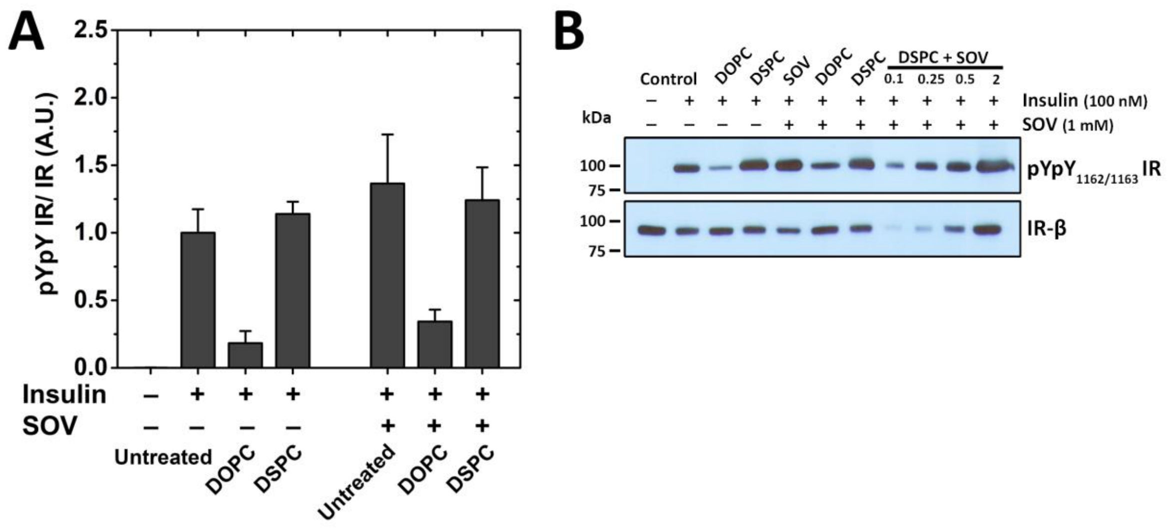


**Supplementary figure 4:** IR activity at 37˚C. (A) Quantification of band intensities using ImageJ from four independent experiments are shown. (B) A representative western blot image is shown.

| **Lipid** | **New PM width in Lo (Å)** | **New PM width in Ld (Å)** |
|---|---|---|
| DLPC | 33.8 | 29.8 |
| DMPC | 35.4 | 31.4 |
| ***DPPC**** | ***37*** | ***33*** |
| DSPC | 38.6 | 34.6 |
| Unexchanged plasma membrane | 37 | 33 |

**Supplementary table 1:** Estimated plasma membrane (PM) domain hydrophobic core width of lipid bilayer after lipid exchange with saturated PCs. Exchange with DPPC (which has 16 carbon acyl chains) is assumed to be similar to untreated normal hydrophobic core width (37 Å for Lo domains and 33 Å for Ld domains from reference [40]). Remaining widths are calculated by adding or subtracting 0.8 Å per carbon acyl chain increase or decrease, based on data in reference [40].

| **Lipid** | **Tm (˚C)** | **Reference** |
|---|---|---|
| bSM | 39 | [55] |
| eSM | 38 | [56] |
| POPC | -4.6 | [57] |
| DOPC | -17.3 | [58] |
| DLPC | -2.1 | [59] |
| DMPC | 23.9 | [59] |
| DPPC | 41.4 | [59] |
| DSPC | 55.3 | [59] |

**Supplementary table 2:** Published gel to Ld phase transition temperatures of single phospholipid vesicles composed of lipids used in this study.

| Structure | CMC in water (mM) | Micelle hydrophobic width (Å) |
|---|---|---|
| n-Decyl-β-D-Maltopyranoside (10 C) 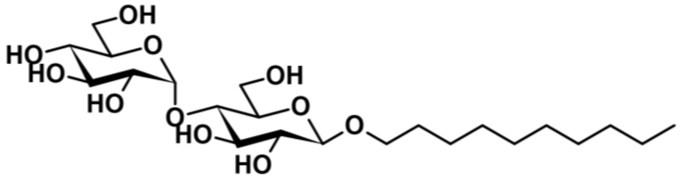 | 1.8 | **27** |
| n-Dodecyl-β-D-Maltopyranoside (12 C) 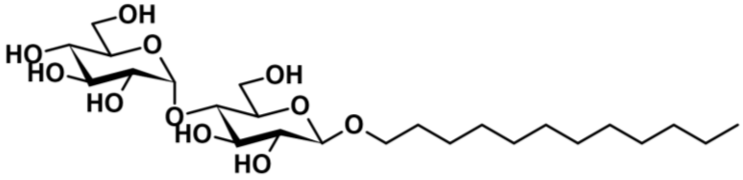 | 0.17 | **32** |
| n-Tetradecyl-β-D-Maltopyranoside (14 C) 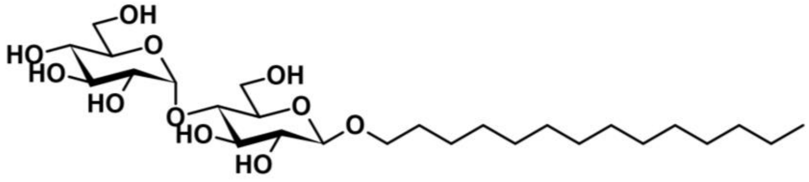 | 0.01 | **37†** |
| n-Hexadecyl-β-D-Maltopyranoside (16 C) 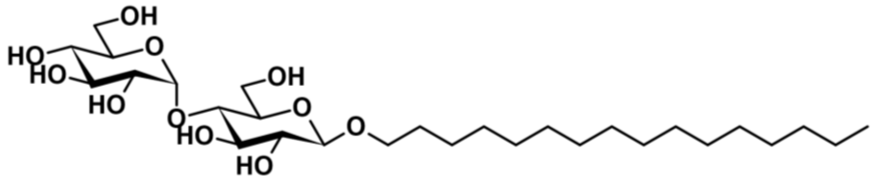 | 0.0006 | **41†** |

†Estimated hydrophobic width.

**Supplementary table 3: Alkyl maltoside detergents used in this study.** Molecular structure of the maltoside detergents used in this study along with their critical micelle concentrations (CMC) and their hydrophobic micelle widths from reference [43] or estimated micelle widths assuming a 4.8 angstrom increase in width as alkyl chain is increased by two carbons
